# Structures and comparison of endogenous 2-oxoglutarate and pyruvate dehydrogenase complexes from bovine kidney

**DOI:** 10.1101/2022.04.06.487412

**Authors:** Shiheng Liu, Xian Xia, James Zhen, Zihang Li, Z. Hong Zhou

## Abstract

The α-keto acid dehydrogenase complex family catalyzes the essential oxidative decarboxylation of α-keto acids to yield acyl-CoA and NADH. Despite performing the same overarching reaction, members of the family have different component structures and structural organization between each other and across phylogenetic species. While native structures of α-keto acid dehydrogenase complexes from bacteria and fungi became available recently, the atomic structure and organization of their mammalian counterparts in their native states remain unknown. Here, we report the cryo electron microscopy (cryoEM) structures of the endogenous cubic 2-oxoglutarate dehydrogenase complex (OGDC) and icosahedral pyruvate dehydrogenase complex (PDC) cores from bovine kidney determined at 3.5 Å and 3.8 Å resolution, respectively. The structures of multiple protein were reconstructed from a single lysate sample, allowing direct structural comparison without the concerns of differences arising from sample preparation and structure determination. Although native and recombinant E2 core scaffold structures are similar, native structures are decorated with their peripheral E1 and E3 subunits. Asymmetric sub-particle reconstructions support heterogeneity in the arrangements of these peripheral subunits. Additionally, despite sharing a similar monomeric fold, OGDC and PDC E2 cores have distinct interdomain and intertrimer interactions, which suggests a means of modulating self-assembly to mitigate heterologous binding between mismatched E2 species. The lipoyl moiety lies near a mobile gatekeeper within the interdomain active site of OGDC E2 and PDC E2. Analysis of the two-fold related intertrimer interface identified secondary structural differences and chemical interactions between icosahedral and cubic geometries of the core. Taken together, our study provides direct structural comparison of OGDC and PDC from the same source and offers new insights into determinants of interdomain interactions and of architecture diversity among α-keto acid dehydrogenase complexes.

## Introduction

The pyruvate dehydrogenase complex (PDC; components suffixed by ‘p’) is an essential element of life that catalyzes the oxidative decarboxylation of pyruvate to acetyl-coenzyme A (acetyl-CoA). This key metabolic reaction links glycolysis to oxidative phosphorylation and the Krebs cycle. PDC is a member of the α-keto acid dehydrogenase complex family, alongside 2-oxoglutarate dehydrogenase complex (OGDC; ‘o’) and branched-chain α-keto acid dehydrogenase complex (BCKDC; ‘b’), which all perform analogous reactions in central metabolism^1,2^. Genomic studies have identified mutations in components of these multienzyme complexes linked to severe clinical consequences, including metabolic acidosis and neurodegeneration^3–7^. PDC also attracts interest in cancer biology for its role in modulating the Warburg effect to promote tumor anabolism^4,8–10^.

PDC, OGDC, and BCKDC are each comprised of multiple copies of a substrate specific dihydrolipoamide acetyltransferase (E2p), dihydrolipoamide succinyltransferase (E2o), or dihydrolipoamide acyltransferase (E2b) inner catalytic (IC) core surrounded by multiple copies of the respective α-keto acid dehydrogenase (E1p, E1o, or E1b) and universal dihydrolipoamide dehydrogenase (E3) components^1,2^. Bovine E2 is comprised of one or two lipoyl domains (LDs) followed by a peripheral subunit-binding domain (PSBD) and an IC domain that are connected by flexible linkers (Fig. 1a). Each α-keto acid dehydrogenase complex proceeds through a three-step mechanism, starting with the decarboxylation of the α-keto acid substrate by E1 and transfer of an acetyl, acyl, or succinyl functional group to lipoate in the LD of E2. Then, the functionalized LD localizes to the IC core, where the functional group is transferred from the lipoyl moiety to CoA. Lastly, E3 reoxidizes the lipoyl moiety, which enables the process to repeat with the production of NADH. The LD translocates between these active sites through a flexible swinging arm mechanism^11^.

**Figure 1.**
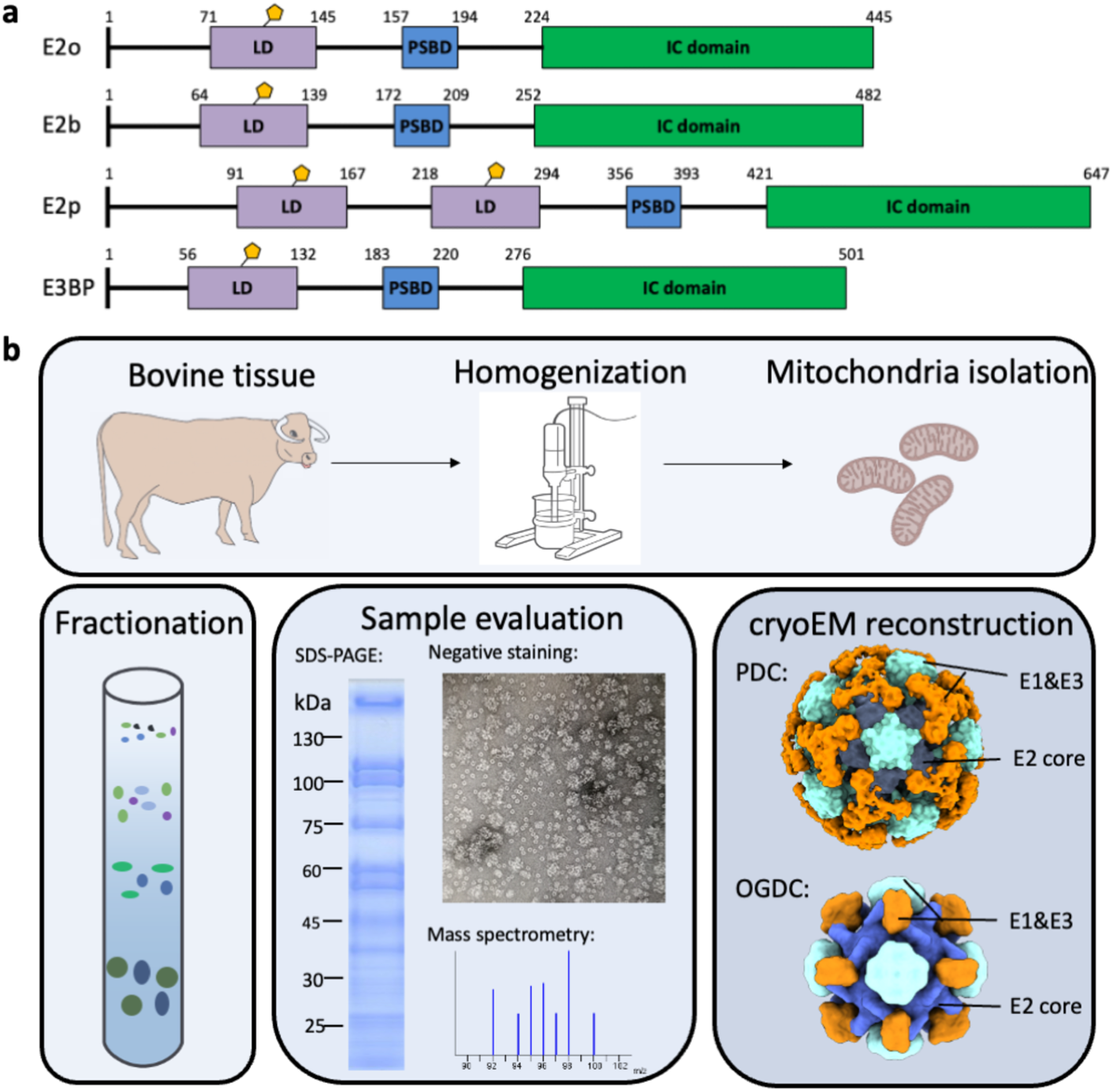
Preparation of bovine OGDC and PDC from native tissue lysate. **(a)** Organization of E2 and E3BP domains. **(b)** Workflow for extraction of PDC and OGDC from bovine tissue lysate. Bovine kidney was homogenized to isolate mitochondria. Mitochondrial lysates were then fractionated by sucrose gradient. Samples were evaluated for α-keto acid dehydrogenase complex by SDS-PAGE, negative stain, and mass spectrometry. CryoEM reconstruction of these samples yielded icosahedral and cubic cores surrounded by peripheral densities.

Although each E1, E2, and E3 subunit in an α-keto acid dehydrogenase is highly similar both in sequence and overall structure to its counterpart in the other classes of the family^12,13^, there are notable differences in the architecture of how these components assemble. E1p is a heterotrimer composed two alpha and two beta subunits, whereas E1o is a homodimer where the alpha and beta subunits are a singular polypeptide^14^. In bacteria, PDC, OGDC, and BCKDC cores all have a 24-mer cubic architecture. As an exception, actinobacteria distinctly lack assembled cores of E2p and possess a unique OdhA protein that fuses the E2o catalytic domain to E1o^15^. However, in eukaryotes, PDC cores adopt a different 60-mer icosahedral architecture despite OGDC and BCKDC cores retaining a cubic geometry^13,16–20^. Additionally, eukaryotic PDC have an additional noncatalytic E3-binding protein (E3BP) that specifically binds E3 to the core and is structurally similar to E2^13,16,21–27^. In mammals, E3BP substitutes for E2p in the core scaffold at possible stoichiometric ratios of 48:12 or 40:20 E2p:E3BP in an unknown arrangement^23,26^. Nonetheless, the subunits of each α-keto acid dehydrogenase complex self-assemble into their respective complexes^28^.

Much of our current structural understanding of α-keto acid dehydrogenase complexes is derived from crystal structures of recombinantly expressed proteins. However, the flexible assembly of intact α-keto acid dehydrogenase complexes is unamenable to crystallography, and components were often crystallized individually. Structures from these individual proteins are unable to fully capture how multiple components come together and function as a complex. Additionally, crystallization may impose artificial order on protein side chains^29^. In the absence of the structure of native complexes, we have an insufficient understanding of how these subunits are organized into functional units. Recently, structures of native PDC from fungus and bacteria became available from cryo electron microscopy (cryoEM)^13,20,27,30,31^. In the intact fungus PDC structures, E3BP was observed as an additional component appended to the interior of the core instead of substituting for an E2p subunit of the core like in mammals and contains a conserved fold found in other E2 proteins^13,27,30^. From bacteria cultured in minimal media to minimize the activity of PDC, cryoEM density of the lipoyl domains interacting with the PDC IC domains could be observed. Despite these exciting progresses, no such endogenous, intact α-keto acid dehydrogenase complexes have been determined for mammals.

In this study, we report the cryoEM structures of endogenous OGDC and PDC E2 IC domain cores extracted from bovine kidney tissue at 3.5 Å and 3.8 Å resolution, respectively. We observe peripheral subunits around the core and the lipoyl moiety substrate within the E2 active site for both complexes. By comparing structures of PDC and OGDC from the same source, we identify distinct interactions in the IC domain trimer and in the two-fold related intertrimer interface, which may direct self-assembly within a milieu of similar components.

## Results

### Extraction, cryoEM, and identification of native, intact OGDC and PDC

To generate samples of intact α-keto acid dehydrogenase complexes, we extracted mitochondrial lysates from bovine (*Bos taurus*) kidney tissue. We then used sucrose gradient fractionation to separate and enrich the protein species present in the mitochondrial lysates (Fig. 1b). Fractions were evaluated by SDS-PAGE, western blot, and mass spectrometry to confirm the presence of PDC and OGDC (Fig. 1b, Supplementary Fig. 1a). The mass spectrometry data indicated BCKDC to be present only as a minor species compared to PDC and OGDC. Fractions containing PDC and OGDC were pooled together into a single lysate sample for EM studies. Negative stain 2D class average evaluation of the sample on a single grid showed both icosahedral classes and cubic classes (Supplementary Fig. S2). Negative stain classes also show peripheral densities around the cores, suggesting that E1 or E3 are present and that intact α-keto acid dehydrogenase complexes were recovered. Higher-order assemblies are visible in cryoEM micrographs (Supplementary Fig. S3a), and smeared densities are present at the periphery of well-resolved cores in the cryoEM 2D classes (Supplementary Fig. S3b). Notably, additional classes belonging to other unknown protein species were obtained (Supplementary Fig. S2 3b), thus highlighting the capability of cryoEM for the study of molecular sociology. CryoEM 3D reconstruction yielded cubic and icosahedral cores surrounded by peripheral densities (Fig. 1b, 2a, 3a).

**Figure 2.**
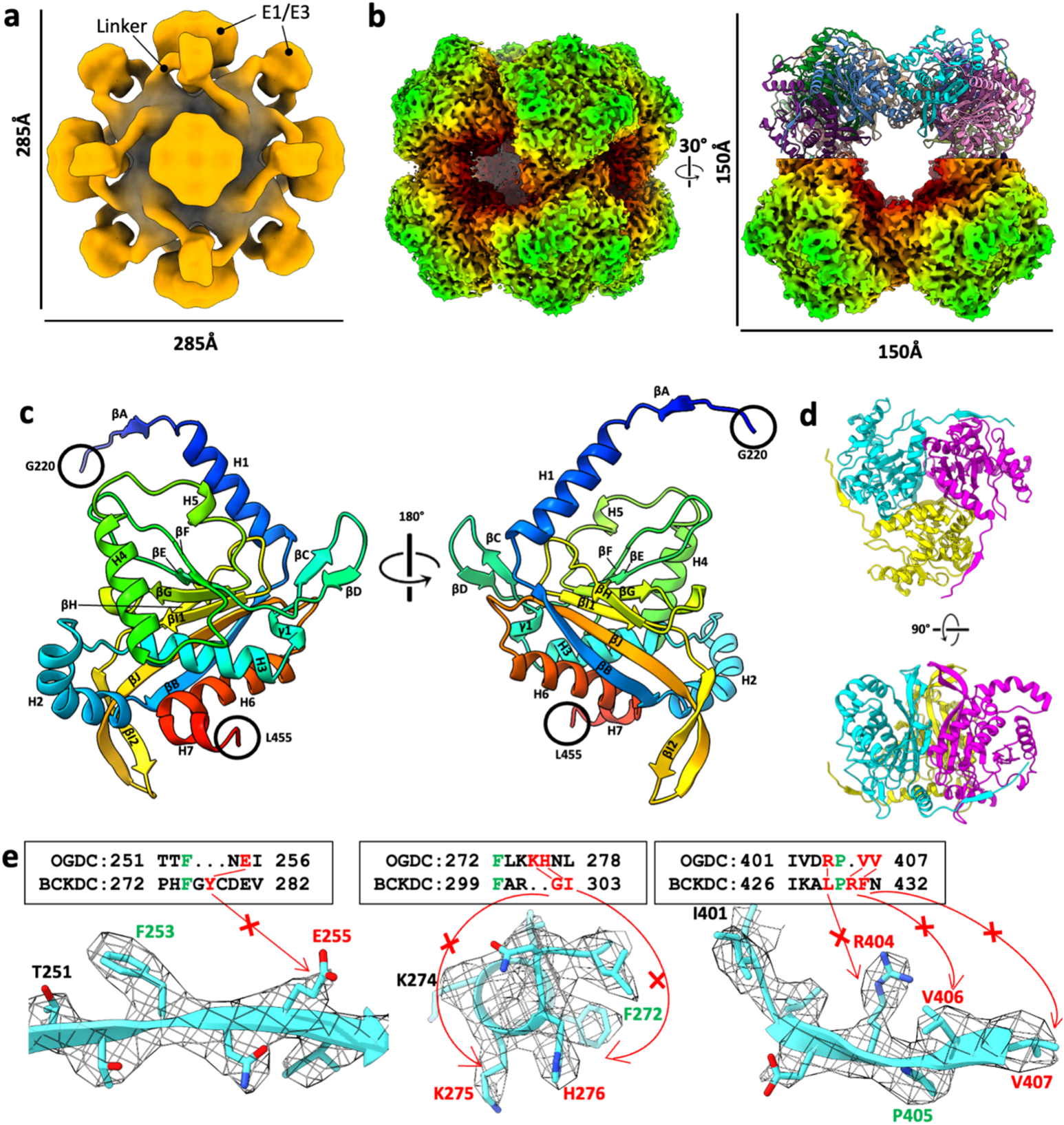
CryoEM reconstruction and identification of bovine OGDC core. **(a)** 3D reconstruction of the full OGDC with octahedral symmetry showing external densities (orange) around the core density (grey). Linkers extend from each E2o subunit towards a density along the edges of the core. Additional large densities are present atop the faces of the core. **(b)** Two views of the cryoEM density map of E2o core colored by radius, with half of our E2o atomic model assembly in the right. **(c)** Rainbow-colored ribbon of E2o IC domain atomic model with secondary structures labeled. Resolvable N-terminal residues (blue) and C-terminus (red) are indicated. **(d)** Structure of the E2o IC domain trimer, colored by subunits. **(e)** Sequence alignment of E2o and E2b with sidechain fit of corresponding E2o sequence into cryoEM density map. Green letters indicate conserved residues between E2o and E2b that establish landmark features in cryoEM density. Red letters and arrows indicate mismatch of residue identity in E2b with corresponding cryoEM density, thus establishing the map to be that of E2o.

**Figure 3.**
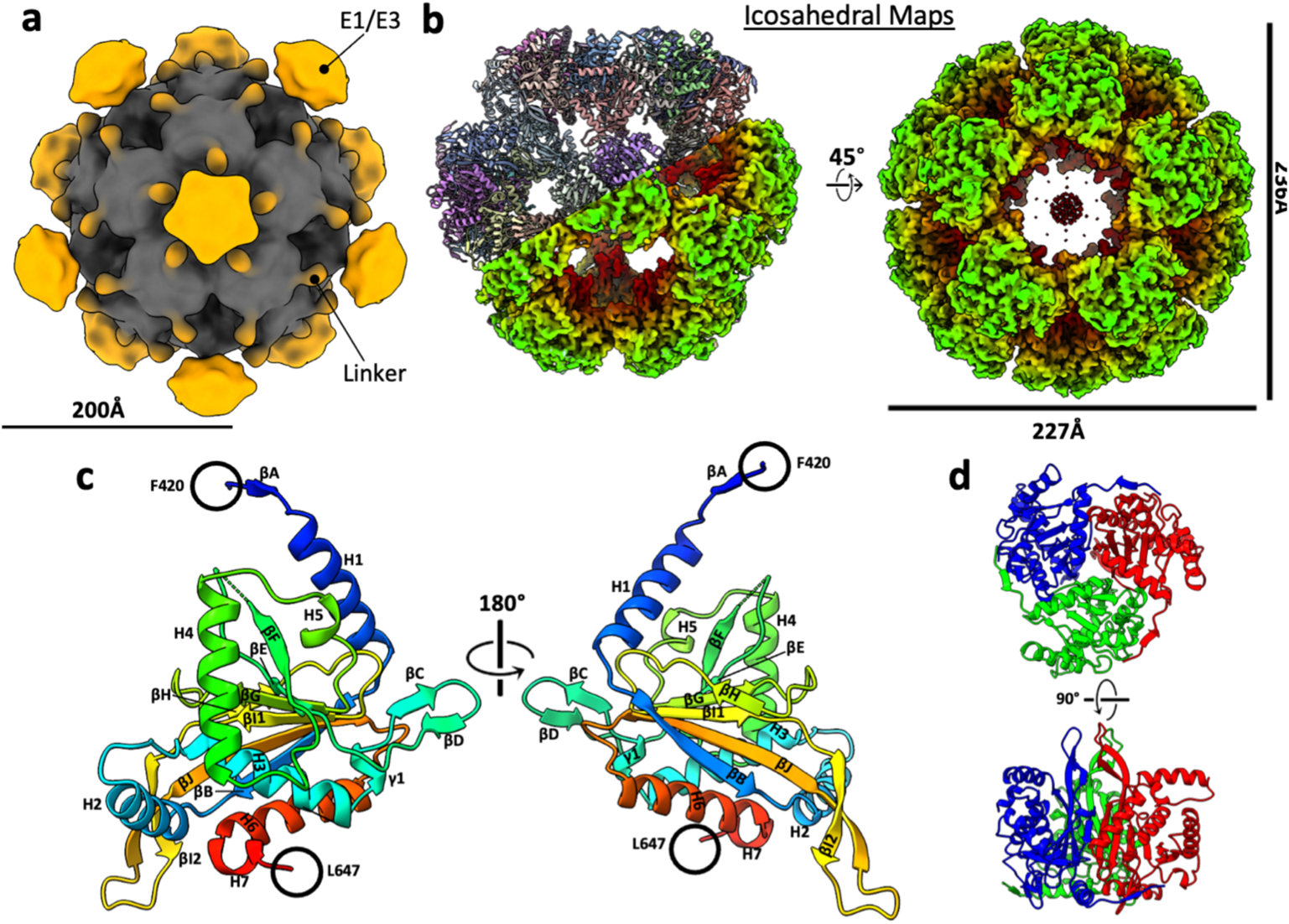
CryoEM reconstructions of bovine PDC and atomic structure of native E2p IC domain. **(a)** 3D reconstruction of the full PDC with icosahedral symmetry shows external densities (orange) around the core density (grey). **(b)** Two views of the cryoEM density map of E2p core colored by radius, with our E2p atomic model assembly of a hemisphere shown in the left panel. **(c)** Rainbow-colored ribbon of E2p IC domain atomic model with secondary structures labeled. Resolvable N-terminal residues (blue) and C-terminus (red) are indicated. **(d)** Two views of the atomic model of the E2p IC domain trimer, colored by subunits.

To structurally characterize the putative OGDC and PDC in these two samples, we obtained 3D reconstructions of the cubic and icosahedral cores of the complexes by single-particle cryoEM at 3.5 Å and 3.8 Å resolution, respectively (Fig. 2b, 3b; Supplementary Fig. S3; Supplementary Table S1). We initially identified PDC based on its distinct icosahedral architecture. This is confirmed by atomic model building of the bovine E2p sequence into the icosahedral cryoEM density map, which showed good agreement between the sequence and the respective side chain densities. FindMySequence results also support the identity assignment of PDC^32^ (E-value 4.80e-74, 3.30e-19, and 2.40e-15 for E2p, E2o, and E2b sequences, respectively).

Although PDC is distinct for its icosahedral architecture, both OGDC and BCKDC share cubic architecture, which introduces ambiguity to the identification of cubic α-keto acid dehydrogenase complexes. To confirm whether our cubic complex was OGDC, BCKDC, or a hybrid of both, we created a homology model by mutating a pre-existing human E2o model to the bovine E2o sequence^19^. After fitting and real-space refinement of the model into our cubic cryoEM density map, the side chains for the E2o model fit the density well (Fig. 2e). Sequence alignment of bovine E2o and E2b reveals primary structure features in E2b that cannot correspond to side chain densities in our map. Fitting of a pre-existing E2b model into our map also showed discrepancies between the modeled side chains and the density^17^. FindMySequence and checkMySequence results also support the identity assignment of OGDC^32,33^ (E-value 5.30e-89 and 6.30e-11 for E2o and E2b sequences, respectively). The unambiguous identification of the cubic complex as OGDC is also reflected in the low presence of BCKDC components in the mass spectrometry data.

### Heterogeneous arrangement of peripheral subunits around the native E2 cores

Unmasked 3D reconstructions have external densities suggestive of E1 or E3 around the core (Fig. 2a, 3a). Both OGDC and PDC have a large density over each core face. In PDC, thin densities from the trimer vertices extend toward the density above the pentameric face. OGDC has additional smaller densities along each cubic edge with thin densities of putative linker regions connecting to the E2o trimer vertices. Previous models based on hydrogen/deuterium exchange and cross-linking mass spectrometry studies propose that E1o binds at the edges and E3 binds at the faces^19,34^. The opposite orientation of the small connecting densities between OGDC and PDC suggests different predominant states of activity and a large range of motion for the flexible linker regions. However, enforced symmetry during reconstruction restricts the interpretability of these external densities for the structures and locations of E1 and E3.

Peripheral subunit densities were further evaluated by symmetry expanded C_1_ sub-particle reconstructions (Supplementary Fig. S4a, b). A 20 Å low pass filter and color zone of 8 Å around the E2 core model were applied to the sub-particles for greater interpretability of low-resolution features (Supplementary Fig. S4c, d). Although the resolution is too low for unambiguous positioning of pre-existing E1 or E3 models within the external densities, we utilized the asymmetric sub-particle reconstructions to narrow down their potential localization. Thin densities corresponding to N-terminal linkers originate from both the two-fold related intertrimer interface and a previously proposed LD binding site in E2o and only the LD binding site in E2p/E3BP^31^. In both PDC and OGDC, extra densities are located near the LD binding sites not adjacent to the face of interest, and fitting of *E. coli* E2p with bound LDs into the E2o sub-particle supports binding of the LD to these sites^31^. In PDC, the extra density at the LD binding site protrudes outward and some have thin extensions (Supplementary Fig. S4d, e). These densities have varying prominence and are notably absent at some IC domains (Supplementary Fig. S4e), suggesting a heterogeneous binding of LDs.

The facial density of OGDC localizes around a single trimer vertex and is connected to the E2o IC domains by the linker originating from the LD binding sites (Supplementary Fig. S4c). Additional smaller densities protrude from the linkers originating from the two-fold related interface. In PDC, only a large facial density is observed (Supplementary Fig. S4d). Similar to that of OGDC, the facial density is localized near a trimer vertex and is connected to the IC domain at the LD binding site. Linkers from the two-fold related interface of E2p/E3BP like those in E2o were not observed. Trajectories of the linker between the IC domain and PSBD from the two-fold related interface and along the three-fold axis have been previously observed but only mutually exclusively^13,15,17–20,25,28,31,35^. In particular, the N-terminal linker of native *E. coli* E2p in the resting state folds back towards the three-fold axis of the trimer after reaching the two-fold related interface^31^. Our asymmetric sub-particle reconstructions suggests that the fold of the E2 N-terminal linker is associated with the activity or reaction step of the complex, and both conformations of the PSBD possibly coexist based on the binding of E1 or E3 (Supplementary Fig. S4c). It remains unknown whether the linker folds back to position the PSBD and LD(s) against its own IC domain or continues past the three-fold axis to a neighboring IC domain.

### E2o and E2p utilize distinct interactions to ensure correct self-assembly into E2 trimers and prevent incorrect heterologous binding of different E2 subunits

For E2o, the backbone of residues 220-455 are fully traceable. For E2p, the backbone of residues 420-647 are fully traceable, except for residues 519 and 520, which lack densities. Both E2o and E2p share conserved secondary structure features: six α-helices (H1-6), a short C-terminal 3_10_-helix (H7), and ten β-strands (βA-J) (Fig. 2c, 3c). The secondary structure elements described here are labeled consistently with *E. coli* E2o, bovine E2b, and human E2p^17,25,36^.

The E2 IC domains of OGDC and PDC are arranged as trimers (Fig. 2d, 3d) with 51 interdomain hydrogen bonds and an average buried surface area of 4957 Å^2^ (standard deviation (SD) = 3.47) buried surface in E2o and an average of 42.75 (SD = 2.36) interdomain hydrogen bonds and an average buried surface area of 4397 Å^2^ (SD = 22) in E2p. Although E2o and E2p share secondary structure features and overall fold, they have notable differences in certain regions and in the interactions that stabilize their respective trimers (Fig. 4). The N-terminus extends at a more elevated angle towards the neighboring IC domain at the clockwise position in E2o than in E2p (Fig. 4a). The N-terminus wraps around a turn between βC′ and βD′ (prime symbol is used to denote the neighboring IC domain at the clockwise position from the exterior view), which is longer in E2o than in E2p. The longer βC-βD turn in E2o enables an electrostatic interaction between the negatively charged surface of the turn and the positively charged surfaces of the N-terminus that is not present in E2p (Fig. 4c). This positions βA in a β-sheet with βD′ and βC′, and the increased stability allowed for modeling of additional N-terminal residues in E2o. The extended βC-βD turn is also present in human E2o and not E2p, but it is lacking in bacteria (Fig. 4b). Additionally, E2p possesses a longer interior hairpin (Fig. 4a), possibly for engaging in interactions in the larger volume of the icosahedral interior of PDC compared to the cubic interior of OGDC^37^.

**Figure 4.**
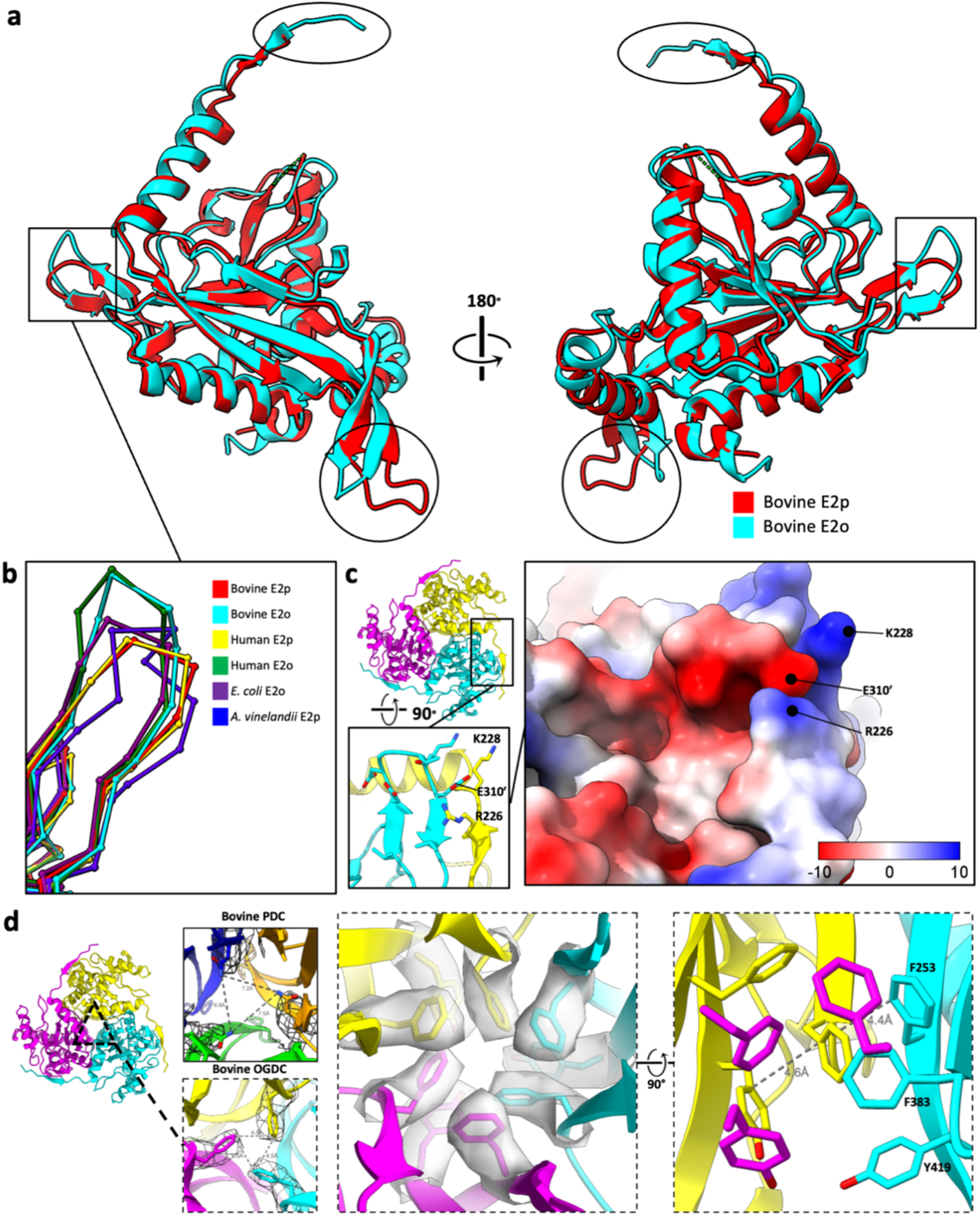
Structure comparison between PDC and OGDC E2 IC domains. **(a)** Superimposition of E2p and E2o IC domains with the N-terminal linker, βC-βD turn, and interior βI2-βJ hairpin indicated by an oval, box, and circle, respectively. E2p has a longer interior hairpin than that of E2o (circle). N-terminal tail diverges at different angles between endogenous recombinant E2o (oval). E2o has a more extended βC-βD turn than that of E2p. (box). **(b)** Comparison of βC-βD turn from E2o and E2p of various organisms (left). **(c)** Location of N-terminal linker and βC-βD turn contact in the E2o trimer. Lower inset shows residue interactions between R226 and K228 of N-terminal linker from the yellow subunit with E310′ of βC-βD turn from the cyan subunit. Right inset: electrostatic surface potential of the same area as that in the lower inset. The positively charged N-terminal linker (primarily blue) associates with the negatively charged βC-βD turn (primarily red). **(d)** π-stacking interactions present at the three-fold axis of E2o trimer but absent in E2p trimer. Expanded views along the three-fold axis of E2o shows an extended π-electron network formed by F253, F383, and Y419.

Within the IC domain trimer, an E2 subunit binds to its clockwise adjacent neighbor through a short β-β motif between βH and βB′. In E2p, this binding is described by three main chain-to-main chain hydrogen bonds (Supplementary Fig. S5). In E2o, this binding is described by two main chain-to-main chain hydrogen bonds, a main chain-to-side chain hydrogen bond, and a slipped-parallel π-stacking interaction^38^ (Phe383-Phe253′). The slipped-parallel π-stacking interaction of E2o occurs near the three-fold axis of the trimer and may form a delocalized π-electron network as previously suggested in *C. thermophilum* E2p between Arg384^30^. By sequence and structure alignment, Phe383 in bovine E2o is the equivalent residue of Arg384 in *C. thermophilum* E2p. Notably, in bovine E2o, the delocalized π-electron network is extended within the three-fold axis and encompasses Phe253, Phe383, and Tyr419 (Fig. 4d). This delocalized π-electron network is also present in human E2o but not in bovine E2p at this structurally analogous position because the equivalent residues Asn575 are too distant from each other^19^ (Fig. 4d). Like mismatched puzzle pieces, the different interfaces of E2o and E2p may prevent incorrect assembly due to co-existence of their components within the cell.

### The lipoyl moiety is positioned near a flexible gatekeeper in the catalytic site

In the cryoEM density maps of E2p, a region of flexibility has been identified: a β-turn connecting βE and βF near the interdomain active site (Fig. 5a). Due to missing density at the βE-βF β-turn, two residues (Ala519 and Gly520) are unmodelled in our E2p model (Fig. 5b). Although the corresponding density is not missing in E2o, it is still weaker relative to that of the adjacent residues. This missing or weak density has also been observed in previous cryoEM density maps^13,15,19,20,31^. The density of the adjacent residue (Leu521 in E2p and Leu329 in E2o) is well-defined. This leucine is a conserved gatekeeper residue for the binding of the lipoyl moiety^17,25^. This gatekeeper residue is proximal to the interdomain active site, near the catalytic residues Ser566 and His620′ in E2p, and positions the dihydrolipoamide (DHLA) into the catalytic site upon binding of CoA, as suggested previously^17^.

**Figure 5.**
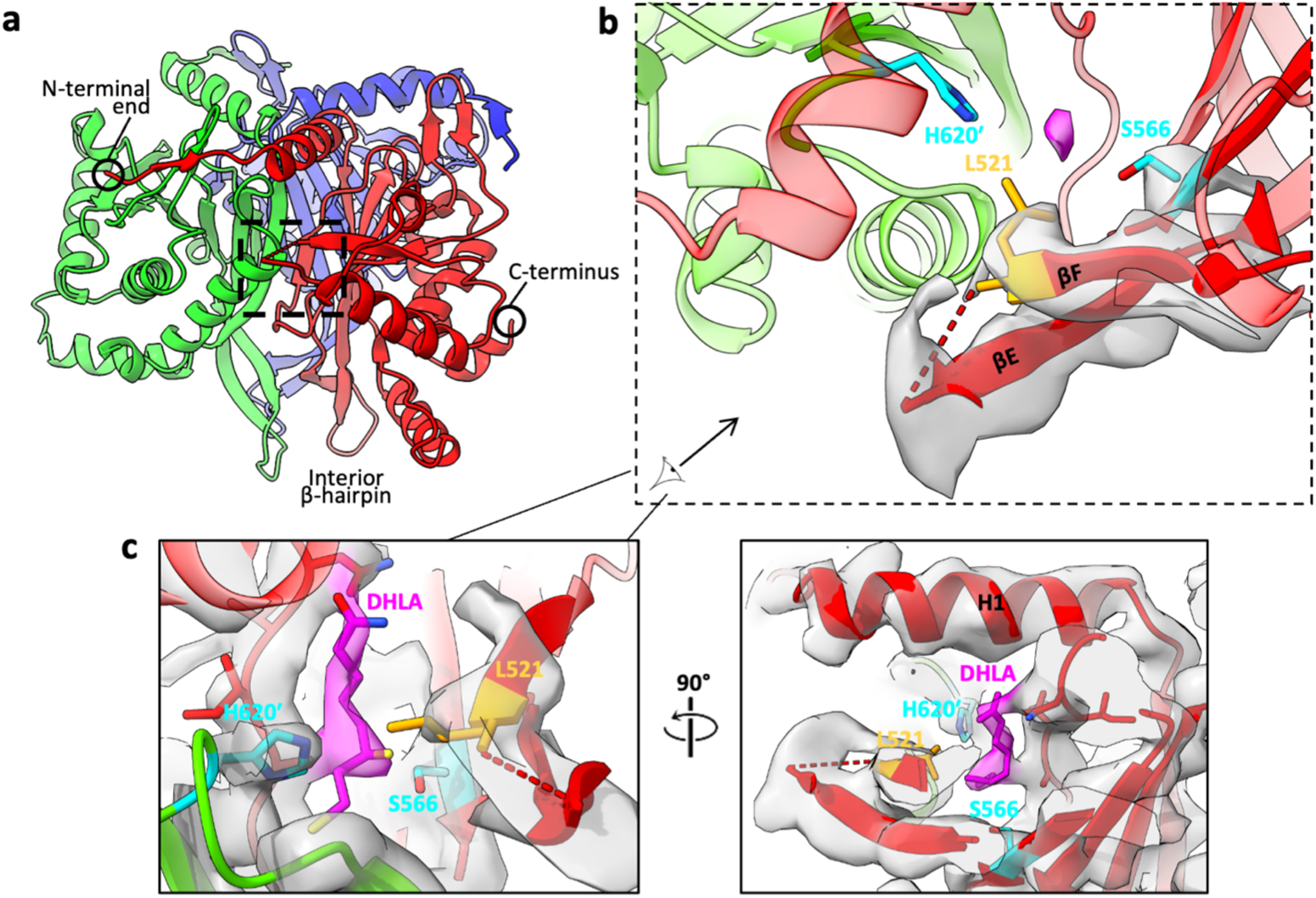
Flexible elements within the bovine PDC E2 IC domain trimers. **(a)** Ribbon-model of E2p trimer (colored by subunits) with boxes highlighting the relative locations of the flexible βE-βF turn (black box). **(b)** Enlarged views of the βE-βF turn near interdomain active site relative to conserved catalytic residues (teal), Ser566 from the red subunit and His620 from the green subunit. Two residues (Ala519 and Gly520) are unmodeled in the turn due to absence of continuous cryoEM density (semi-transparent gray; for clarity, only those for βE-turn-βF are shown), suggesting flexibility of the residues in the turn. The conserved leucine gatekeeper (orange) is next to these unmodeled residues. A weak density (pink) bordered by catalytic and gatekeeper residues within active site is visible. **(c)** CryoEM density of the same region of (B) shown at a decreased threshold as semi-transparent pink for the lipoyl moiety or gray for the surrounding region. Atomic models for the lipoyl moiety (from PDB ID: 1EAE) and the surrounding area are shown as sticks and ribbon, respectively.

In the E2p cryoEM density map, a putative density for the lipoyl moiety was identified near the catalytic residues Ser566 and His620′ and the gatekeeper residue Leu521 (Fig. 5b). When the contour level is decreased, the density forms a thin extension through the active site channel and accommodates a model of DHLA^35^ (adopted from PDB ID: 1EAE) (Fig. 5c). The E2o cryoEM density map also contains a putative lipoyl moiety density (Supplementary Fig. S6a). Following the trajectory of these densities, the dihydrolipoyllysine enters the active site from the exterior of the E2 core at an entrance at the end of a cleft between H1 and a turn connecting βH and βI1, as previously depicted in *E. Coli* E2p in the resting state^31^. Densities of the lipoyl moiety are not present in cryoEM density maps of native *C. thermophilum* E2p and E2o and *N. crassa* E2p^13,20,30^. In the E2o density map at low contour level, weak cylindrical densities reminiscent of β-strands from the LD or helices from the PSBD are observed to the left and to the right of the entrance of the cleft. The left densities are located similarly to that of the LD in native *E. coli* E2p in the resting state^31^. (Supplementary Fig. S6b). These additional densities extend along the cleft towards the active site entrance. We attempted to fit an LD model and PBSD models from NMR (PDB ID: 1LAC; 1W4H) and Alphafold prediction^39–42^ (AF-P11179-F1; AF-P11181-F1), but there are mismatches between the cylindrical densities and the LD models when the dihydrolipoyllysine-containing loop is positioned adjacent to the active site channel. These observations combined with structural variation between LD domain models suggest that the β-sheets of the LD may be conformationally variable and undergo conformational and positional changes during catalysis.

To search for E3BP, we expanded the symmetry of the trimer vertex and pentameric face sub-particles and reconstructed them asymmetrically (C_1_) (Supplementary Fig. S7a). Due to low resolution of the C_1_ reconstructions, we are unable to utilize side chain densities to distinguish sequence identity. Sequence alignment between E2p and E3BP and fitting of predicted E3BP structures (PDB ID: 6H60; AF-P22439-F1) show two notable regions of difference in E3BP within the IC domain^26,40,41^: a three-residue shorter linker between H2 and H3 and a three-residue longer interior hairpin between βI2 and βJ (Supplementary Fig. S7b). We searched the classes of the trimer vertex and pentameric C_1_ sub-particles but were unable to identify any differences corresponding to the presence of E3BP (Supplementary Fig. S7c). E3BP is not evenly distributed in the trimers^26^, and current 3D classification methods may place too much weight on the larger features in the conserved fold of E2p and E3BP, which may reduce the quality of asymmetric reconstructions of the interior hairpin. Heterogeneity of E3BP distribution precludes identification and experimental structural determination of E3BP from native mammalian PDC.

### Different knob-and-socket intertrimer interactions prevent heterologous self-assembly of E2 trimers

E2o and E2p have near-identical structures, yet they assemble into core scaffolds with different geometry. The cubic and icosahedral geometries are related by principles of quasi-equivalence and Euclidean geometry^28^. Each IC domain trimer vertex of the core is bound to its neighboring trimers through a palindromic two-fold related intertrimer interface^19,25^ (Fig. 6a; Supplementary Fig. S8a). The hydrophobic C-terminal 3_10_-helix (H7) of each IC domain binds to a hydrophobic pocket formed by residues of H2*, H7*, and the N-terminal end of H4* (asterisk is used to denote the partner IC domain across the two-fold related interface) on the opposite IC domain in a knob-and-socket interaction (Supplementary Fig. S8b). This binding is further stabilized by electrostatic interactions from oppositely charged surfaces lining the exteriors of H7 and the hydrophobic pocket (Supplementary Fig. S8c). In actinobacteria E2p, where H7 folds back against rather than away from the IC domain, the knob-and-socket interaction cannot occur, so the trimers are unable assemble into a core^15^.

**Figure 6.**
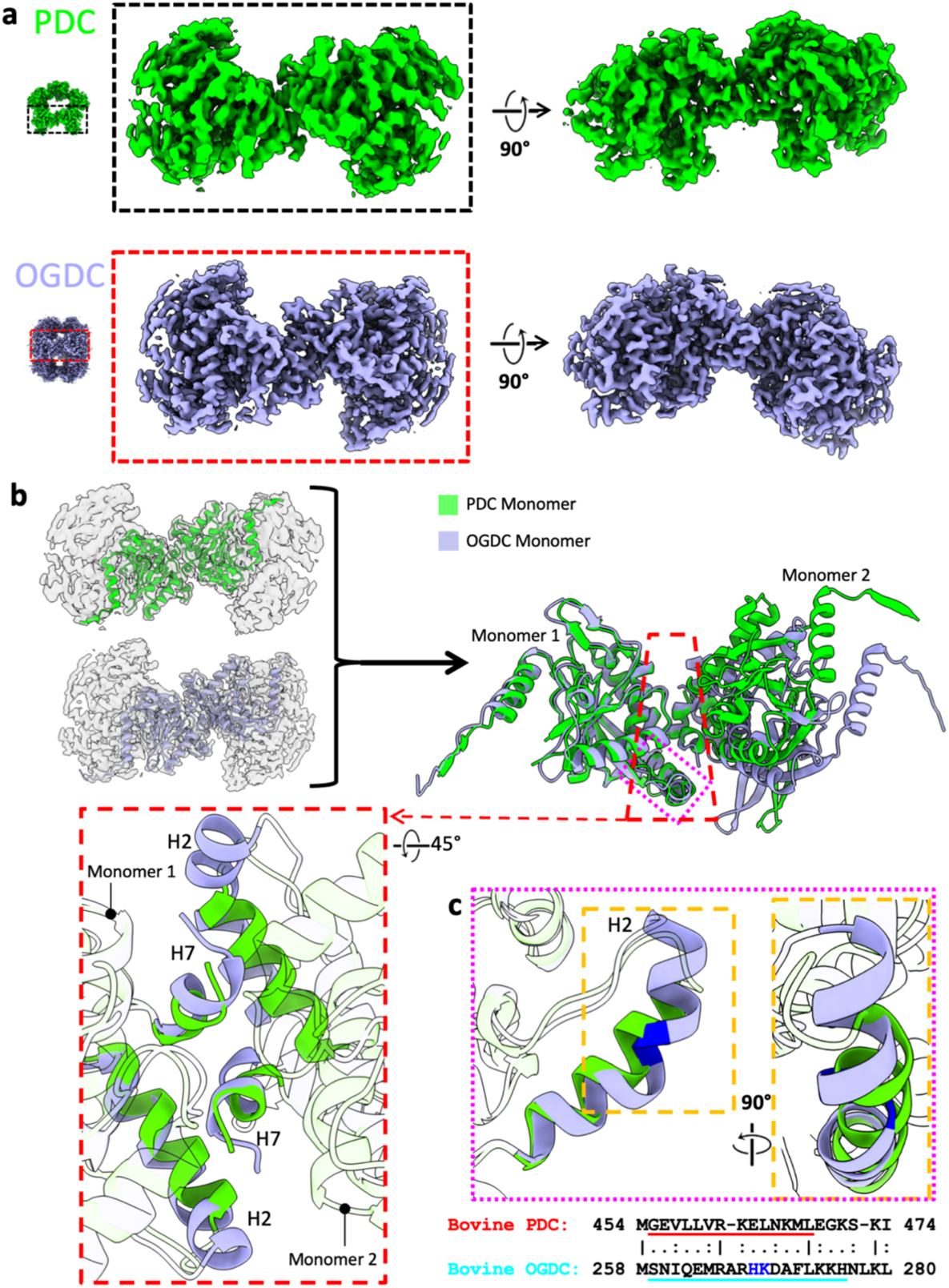
Differences in the E2 inter-trimer interactions between PDC and OGDC. **(a)** Two orthogonal views (perpendicular and parallel to the 2-fold axis) of interacting E2 trimers from the cryoEM density maps of PDC (green) and OGDC (blue). **(b)** Comparison of the intertrimer interface in PDC (green) and OGDC (blue). Interactions involve only two subunits, one from each trimer, as apparent from the left panel where the atomic model (ribbons) of the interacting subunits are superposed in their respective densities. The two interacting subunits shown in (A) are aligned through their left subunit (monomer 1) and shown together to identify differences at the interacting interfaces with boxed regions shown as insets to highlight differences of their corresponding H2 helix. The inset in red box shows the knob-and-socket interaction of H2 and H7 (colored; rest of ribbon transparent for clarity) at two-fold interface in E2p and E2o. **(c)** Superimposition and sequence alignment of H2 in E2p and E2o. The helix-kink-helix (kink highlighted in blue) is present in E2o but absent in E2p. Inset in the orange box: Enlarged orthogonal view of the diverging H2 trajectories.

The symmetries for bovine E2o and E2p are obtained from a different combination of interactions (Supplementary Table S2). The intertrimer interface has an average of 18.7 (SD = 0.49) hydrogen bonds and an average buried surface area of 1102 Å^2^ (SD = 0.46) in E2o and an average of 5.6 (SD = 0.63) hydrogen bonds and an average buried surface area of 825 Å^2^ (SD = 14.18) in E2p. The exterior of the E2o socket is more charged than that of E2p (Supplementary Fig. S8c), which leads to the greater number of hydrogen bonds at the E2o intertrimer interface. Additionally, the exterior surface of the socket in E2o and E2p are oppositely charged from one another. Although the residues comprising the hydrophobic sockets between the two species are similar, the shapes of the sockets are different to accommodate likewise different knobs. The E2p knob has one hydrophobic surface that binds a single hydrophobic pocket in the socket. However, the E2o knob has an additional hydrophobic surface on the opposite side of the knob, and the E2o socket contains two pockets to accommodate this difference (Supplementary Fig. S8b). Similar to the purpose of the interdomain interface differences to prevent heterologous assembly of E2 subunits, the oppositely charged exterior surfaces and differently shaped knob-and-socket of the intertrimer interface between E2o and E2p suggest a means of preventing heterologous assembly of E2 trimers during core formation.

To compare the relative positions of trimers at the intertrimer interface between E2o and E2p, the IC domains of the two-fold related interface were aligned by superimposition at one trimer vertex (Fig. 6b). The fold of H2 and the position of H7 differ between E2o and E2p. H7 of E2o (Pro448-Leu453) has a length of six residues and has two additional residues (Asp454 and Leu455) appended at the C-terminus. H7 of E2p (Pro642-Leu646) has a length of five residues and one additional residue (Leu647) at the C-terminus. H7 is positioned more closely to H4* in E2p than in E2o, which is enabled in E2p by an additional hydrogen bond between H7 and H4* and fewer hydrogen bonds between H7 and H2*. (Supplementary Table S2). In E2o, H2 has a length of 18 residues (Ser259-His276) and is comprised of a helix-kink-helix with the kink starting at residue 268. In E2p, H2 has a length of 14 residues (Gly455-Leu468) and is a single, uninterrupted helix (Fig. 6c). The H2 kink in E2o positions the C-terminal end of the helix more closely against H7* for hydrogen bonds between Lys275 and His276 with Asp454* and Asp454*, respectively. The different interactions at the knob-and-socket stabilize E2 IC trimers at angles that enable the Euclidean geometry relationship between a cube and icosahedron.

## Discussion

Advances in cryoEM and its integration with mass spectrometry analysis have enabled the structural characterization of intact, native complexes^32,43,44^. Endogenous methods have recently been applied to determine structures of PDC in fungi and bacteria^13,20,27,30,31^. Here, we extended these methods to the study of mammalian tissues and determined structures of native bovine OGDC and PDC at sufficient resolutions for model building of their E2 IC domains. Analysis of OGDC and PDC from the same source enables direct comparison between the two protein species. Although E2 IC domains share a nearly identical fold to conduct analogous reactions, nuanced differences in their structures highlight the divergent evolution of α-keto acid dehydrogenase complexes to meet the requirements of different organisms with varied cellular contents and metabolic environments.

Besides the structures reported here, native structures of α-keto acid dehydrogenase complexes from the same endogenous source only became available early this year for the fungus *C. thermophilum*, where both PDC and OGDC were structurally characterized^20^. In contrast to our bovine E2o and E2p structures, the back-folded N-terminal region comprised of βA and H1 is absent in *C. thermophilum* E2p. Interestingly, H1 is still present in *C. thermophilum* E2o from the same size-exclusion chromatography fraction^20^, and is also present in native *N. crassa* and *E. coli* E2p^13,31^. Given that the structured conformation of the N-terminal region is stabilized by electrostatic interactions between βA and H1 with the βC-βD turn (Fig. 4c), the back-folded conformation may exist transiently depending on the activity state of the complex.

The unfolded state has been suggested to be necessary for unhindered insertion of the lipoyl moiety^30^. For *E. coli* E2p in the absence of pyruvate, H1 is folded and binding of the N-terminus extends towards the three-fold axis, which was suggested to position and stabilize the E2-LD complex in a resting state^31^. Our sub-particle reconstructions show the N-terminal region extending outwards from both the two-fold related interface and about the three-fold axis simultaneously (Supplementary Fig. S4c), which indicates that multiple conformations of the PSBD linker are present. The conformation of the linker may correspond to the transient binding of E1 and E3. There could be a potential mixture of activity or assembly states in different lysate fractions when complexes are sorted by size, and our single-particle cryoEM analysis of the entire core might have only reconstructed the most stable or populous conformation or a potential hybrid of the active and resting states.

The interdomain interactions between an E2 IC domain and its neighbor are not conserved between E2p and E2o. Bovine and human E2o contain an extensive delocalized π-electron network at the three-fold axis (Fig. 4d) that is absent in their E2p counterparts^19,25^. The differences in interdomain interfaces between E2p and E2o within an organism may enable correct self-assembly within a milieu of components in the mitochondrial matrix. *C. thermophilum* E2o may also have a similar delocalized π-electron network, but instead with Met350 at the analogous position of Phe383 in E2o^20^. Met-aromatic motifs can provide additional stabilization compared to purely hydrophobic interactions^45^. Rather than a delocalized π-electron network of aromatic residues, *C. thermophilum* E2p instead possesses a potential stabilizing arginine cluster that is absent in both its E2o counterpart, mammalian E2p, and mesophilic fungus *N. crassa* E2p^13,20,30^. The greater interdomain interactions could contribute towards increased thermal stability needed for thermophiles compared to mesophiles as a result of divergent evolution.

While there are pre-existing structures of α-keto acid dehydrogenase complexes, the diversity of interactions within these complexes between different organisms occludes direct comparison for determinants of assembly. Although *E.coli* E2p and bovine E2b are both cubic like bovine E2o, their interdomain interface interactions differ greatly. The knob of *E. coli* E2p has a semicylindrical hydrophobic surface that complements a narrow, elongated hydrophobic socket^31^. Bovine E2b has a double-sided knob like bovine E2o^17^, but the hydrophobic surfaces are rotated about and have different sizes. Although α-keto acid dehydrogenase complexes can share the same geometry, the interdomain interactions that hold the complex cores together differ between both family members and model organisms. Thus, structural characterization of samples from the same organism are needed to describe distinct interactions.

In mammals, E3BP substitutes for a corresponding E2 in the core scaffold as determined by biochemical analysis instead of being an additional subunit as it is in fungus^13,23,26,27^, and no internal density was observed for our native bovine PDC structure. Because of the heterogeneous arrangement of E3BP and peripheral E1 and E3 subunits, PDC and other α-keto acid dehydrogenase complex may need to be studied at the individual level. Tomography could identify peripheral subunits and avoid issues of classification for E3BP if sufficient resolution could be reached in the future.

Advancements in cryoEM have enabled atomic modeling of native α-keto acid dehydrogenase complexes and other endogenous complexes. Looking forward, faster direct electron detectors and improved cryoEM grid preparation methods will improve acquisition of particles for low population species for unambiguous identification and atomic modeling. Heterogeneous reconstruction could enable visualization of distinct protein conformations^46,47^. Machine-learning based methods in particle picking^48^, structure prediction^40^, and identification can resolve unknown identities in a society of proteins^32^. Structural determination of native geometrically variable complexes from similar folds, exemplified by the cubic and icosahedral complexes presented here, should not only provide new biological insight but also inform protein engineering applications by providing an opportunity for comparison within the same environment to identify the determinants of protein architecture.

## Materials and methods

### Preparation of bovine mitochondria

Bovine mitochondria were prepared from *Bos taurus* kidneys as described with modifications^49^. 6 kg of bovine kidneys were collected and chilled on ice immediately after slaughter. Cortical tissues were cut into small slices (0.5-1 cm) and soaked in 4 L water for 1 h and then washed with 2 L <u>kidney buffer</u> (20 mM potassium phosphate pH 7.6, 250 mM sucrose, 1 mM EDTA). The slices were passed through an electric meat grinder, and the ground meat was suspended in kidney buffer with 10 mM 2-mercaptoethanol and diluted to a final volume of 12 L. The suspension was homogenized using an Ultra-Turrax (IKA) homogenizer (40 s), filtered through 8 layers of cheesecloth accompanied with sieve, and diluted to a total volume of 18 L using kidney buffer. The resulting supernatant was centrifuged at 2,000-3,000 × g for 10 min. The supernatant was decanted and further centrifuged at 5,600 × g for 25 min. The lysosome-enriched fluffy layer was carefully removed from the mitochondrial pellet. The pellets were resuspended in a total of 8 L kidney buffer and filtered again through 8 layers of cheesecloth. The suspension was homogenized using an Ultra-Turrax homogenizer (30 s), diluted to 12 L with kidney buffer, and centrifuged at 5,600 × g for 25 min. The pellets were resuspended with kidney buffer with 0.2 mM PMSF, 0.25 μg/ml aprotinin, 0.14 μg/ml pepstatin, and 1 μg/ml leupeptin in a total volume of 4 L. The suspension was treated with 0.01% digitonin for 15 min to remove outer mitochondrial membranes and diluted to 12 L using kidney buffer. The resulting mitoplasts were concentrated by centrifugation at ~25,600 × g for 20 min followed by removal of excess kidney buffer. The pellet of the final step was flash frozen in liquid nitrogen and stored at −80 °C.

### Isolation of bovine PDC and OGDC complexes

100 g of frozen mitoplasts were thawed in 150 mL lysis buffer [20 mM potassium phosphate pH 7.6, 50 mM NaCl, 0.02 mM thiamin pyrophosphate, 20 mM MgCl_2_, 2 mM EGTA, protease inhibitors (2 mM PMSF, 2.5 μg/mL aprotinin, 1.4 μg/mL pepstatin, and 10 μg/mL leupeptin)]. The sample was diluted to 337.5 mL with lysis buffer. 37.5 mL <u>Triton X-100 buffer</u> [lysis buffer components in addition with 16% (v/v) Triton X-100] was then added, and the sample was stirred for 15 min at 4 °C. The suspension was centrifuged (SLA-3000, 13,000 rpm, 20 min, 4 °C), and the supernatant was PEG precipitated in 5% (w/v) PEG 10,000 for 15 min. The precipitate was collected by centrifugation (2,500 × g, 7 min, 4 °C) and re-suspended in 35 mL lysis buffer. The suspension was homogenized using a Dounce homogenizer and centrifuged (SW28 Ti rotor, 28,000 rpm, 17 min, 4 °C). The resulting supernatant was loaded onto 50% (w/v) sucrose cushions (15 ml) and centrifuged (SW41 rotor, 40,000 rpm, 24 h, 4 °C). Pellets were each dissolved in 0.5 mL sample buffer (50 mM MOPS-KOH buffer pH 7.0, 0.02 mM thiamin pyrophosphate, 10 mM MgCl_2_, 0.1 mM EGTA, protease inhibitors, 1mM DTT) by shaking at 230 rpm for 1 h and clarified by centrifugation (SLA-3000, 16,000 rpm, 20 min, 4 °C). The sample was loaded onto 10–40% (w/v) sucrose gradients (1 mL per gradient) and centrifuged (SW41 Ti, 26,000 rpm, 12 h, 4 °C). The gradients were fractionated and assessed by SDS-PAGE and western blot using a PDC complex antibody cocktail (Abcam) that contains four different mAbs reacting specifically with E1α, E1β, and E2/E3BP. The fractions containing PDC and OGDC complexes were pooled, dialyzed to <u>final buffer</u> (50 mM MOPS-KOH buffer pH 7.0, 0.02 mM thiamin pyrophosphate, 1 mM MgCl_2_, 1 mM NAD+, 0.1 mM EGTA, protease inhibitors, 1 mM DTT), and concentrated for cryoEM analysis. Liquid chromatography-tandem mass spectrometry was performed on the sucrose gradient fractions. Gradient fractions corresponding to OGDC were subjected to gel filtration on a Superose 6 column (GE Healthcare) to remove ferritin.

### CryoEM sample preparation and image acquisition

CryoEM grids were prepared by using an FEI Mark IV Vitrobot (Thermo Fisher Scientific). 3 μL of sample was applied onto a glow-discharged lacey carbon copper grid with a thin, continuous carbon film (Ted Pella). After waiting for 30 s, the grid was blotted (8°C, 100% humidity, 10 s blotting time, 1 blotting force) and plunge-frozen in liquid ethane. Grids were loaded into a Titan Krios (Thermo Fisher Scientific) equipped with a Gatan Imaging Filter (GIF) Quantum LS and a Gatan K2 Summit direct electron detector. Movies were recorded with SerialEM at super-resolution mode^50^. The nominal magnification was 105,000 ×, corresponding to a calibrated pixel size of 0.68 Å at the specimen level. The total exposure time for each movie was set to 8 s and fractionated equally into 40 frames, leading to a total dosage of ~45 electrons/Å^2^. Defocus was set to −1.8 to −2.6 μm.

### Image processing

The movies were motion corrected with MotionCor2 (Zheng, 2017). 6,959 good images were selected from a total of 9,029 by manual screening. After defocus determination by CTFFIND4^51^, particles were automatically picked using Gautomatch (https://www2.mrc-lmb.cam.ac.uk/research/locally-developed-software/zhang-software/). 33,138 particles for PDC complex and 41,165 particles for OGDC complex were obtained by 2D and 3D classification in RELION-3^52^. FSC calculations were performed in RELION-3.

For PDC complex, all particles from the previous step were recentered and extracted with a box size of 576 pixels. These particles were refined with icosahedral symmetry, resulting in a map with a resolution of 3.8 Å. To get better density for the peripheral E1 and E3 subunits, a structure-guided sub-particle strategy was used. First, the STAR file from refinement was expanded with icosahedral symmetry, leading to a total of 1,988,280 particles. Second, the center of the expanded particles was shifted to one trimer or pentameric face. Finally, 3D classification and refinement focused on the putative E1 and E3 region was performed.

For OGDC complex, all particles from the previous step were recentered and extracted with a box size of 384 pixels. These particles were refined with octahedral symmetry and a mask around the E2 core, resulting in a map with a resolution of 3.5 Å. To get better density for the peripheral E1 and E3 subunits, a structure-guided sub-particle strategy was used. First, the STAR file from refinement was expanded with octahedral symmetry, leading to a total of 987,960 particles. Second, the center of the expanded particles was shifted to one face of the cubic core, and particles were extracted in RELION with a box size of 208. Finally, 3D classification and refinement of the external area was performed.

### Atomic modeling

For model building, human E2o (PDB ID: 6H05) and human E2p (PDB ID: 6CT0) were used as templates for bovine E2o and E2p IC domains, respectively^19,25^. Each monomeric subunit model was fitted into the respective cryoEM density map of the core scaffold using ChimeraX^53^, mutated and manually refined in Coot^54^, and real-space refined in Phenix^55^. The monomer model was then duplicated to the appropriate stoichiometry and fit into the respective map and refined iteratively using ISOLDE and Phenix^56^. FindMySequence and checkMySequence were later used to confirm the identity of OGDC^32,33^. Electrostatic potential maps were calculated in ChimeraX. Sequence alignments were performed [Sequence alignments performed with Clustal Omega and EMBOSS Needle on the EMBL-EBI server^57–59^. Structure predictions were obtained from Alphafold using Colabfold or the EBI database^40,41,60^.

### Data availability

Mass spectrometry data and source gel/blot images are provided with this paper. CryoEM density maps of the icosahedral bovine PDC core and cubic bovine OGDC core have been deposited in the Electron Microscopy Data Bank under accession numbers EMD-XXXX and EMD-XXXX, respectively. The coordinates of the bovine E2p IC domain and E2o IC domain have been deposited in the Protein Data Bank under accession numbers XXXX and XXXX, respectively.

## Acknowledgments

This research was supported in part by grants from NSF (DMR-1548924) and NIH (R01GM071940). J.Z. acknowledges support from an Interdisciplinary Training in Virology and Gene Therapy training grant (NIH 5T32AI060567). We acknowledge the use of resources in the Electron Imaging Center for Nanomachines supported by UCLA and grants from NIH (S10RR23057, S10OD018111, and U24GM116792) and NSF (DBI-1338135). We thank the UCLA Proteome Research Center for assistance in mass spectrometry. **Contributions** Z.H.Z. conceived the project and supervised the research. S.L designed the experimental protocols. S.L. and X.X. prepared samples, collected cryoEM images, and determined the 3D structures. Z.L. and J.Z. built atomic models and made figures. J.Z. interpreted the results and wrote the manuscript; all authors reviewed and approved the paper.

---

**Supplementary Fig. 1.**
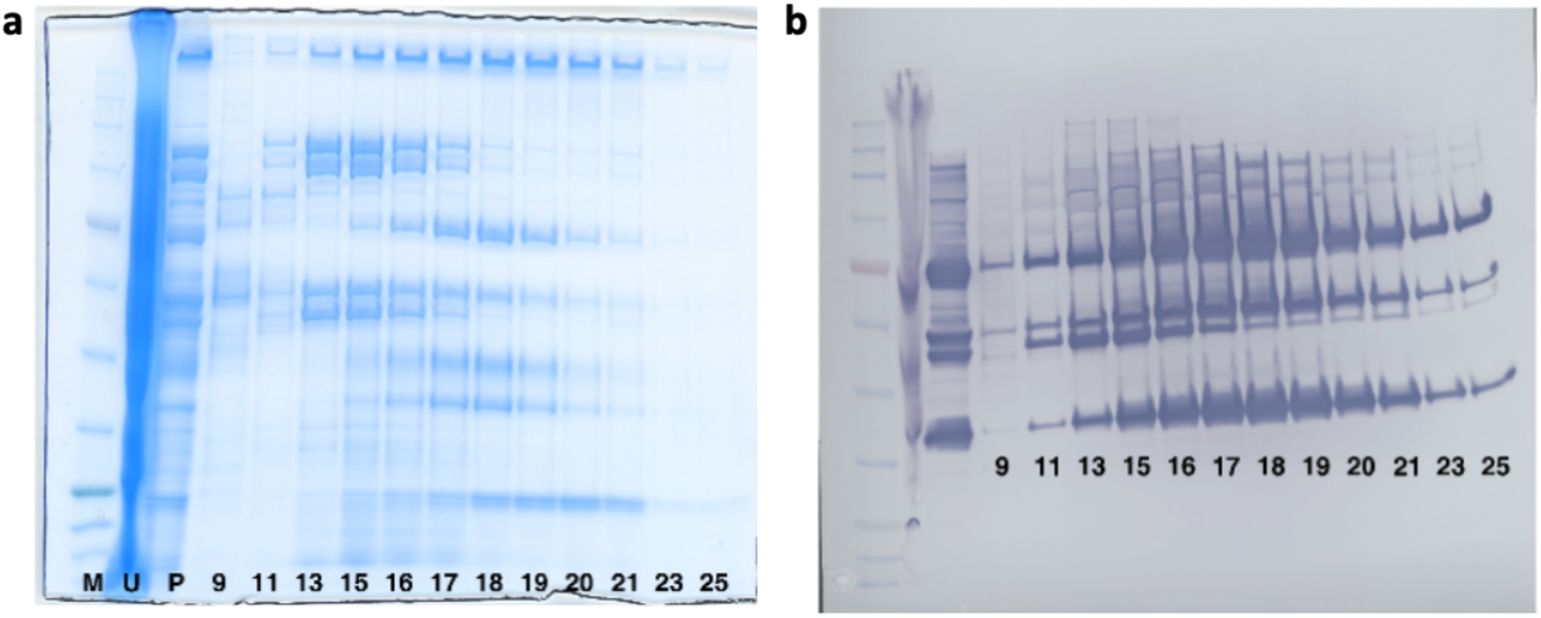
Raw images of gels and blots. **(a)** SDS-PAGE. **(b)** Immunoblot. Numbers below lanes indicate sucrose gradient fraction.

**Supplementary Fig. 2.**
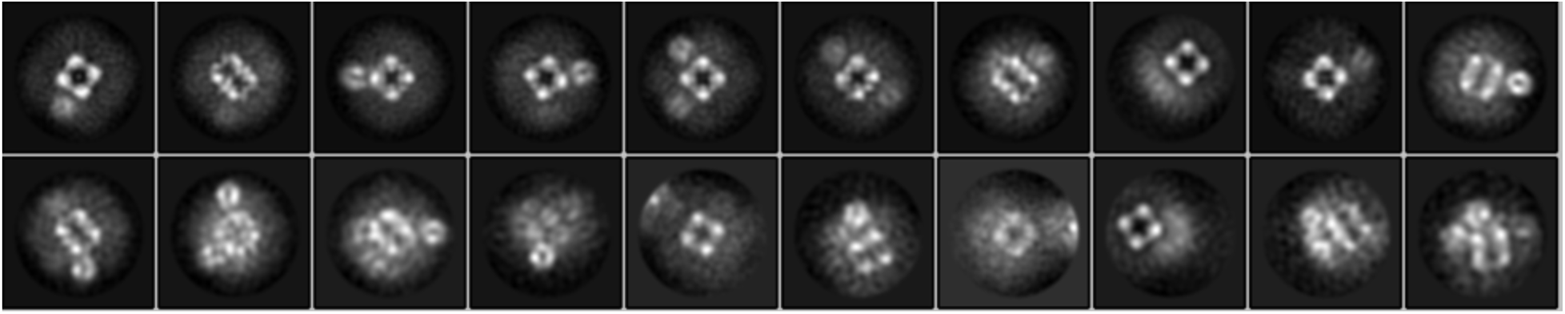
Negative stain 2D class averages. Negative stain 2D class averages show cubic and icosahedral cores with peripheral density. Additional classes of unknown protein species were also observed.

**Supplementary Fig. 3.**
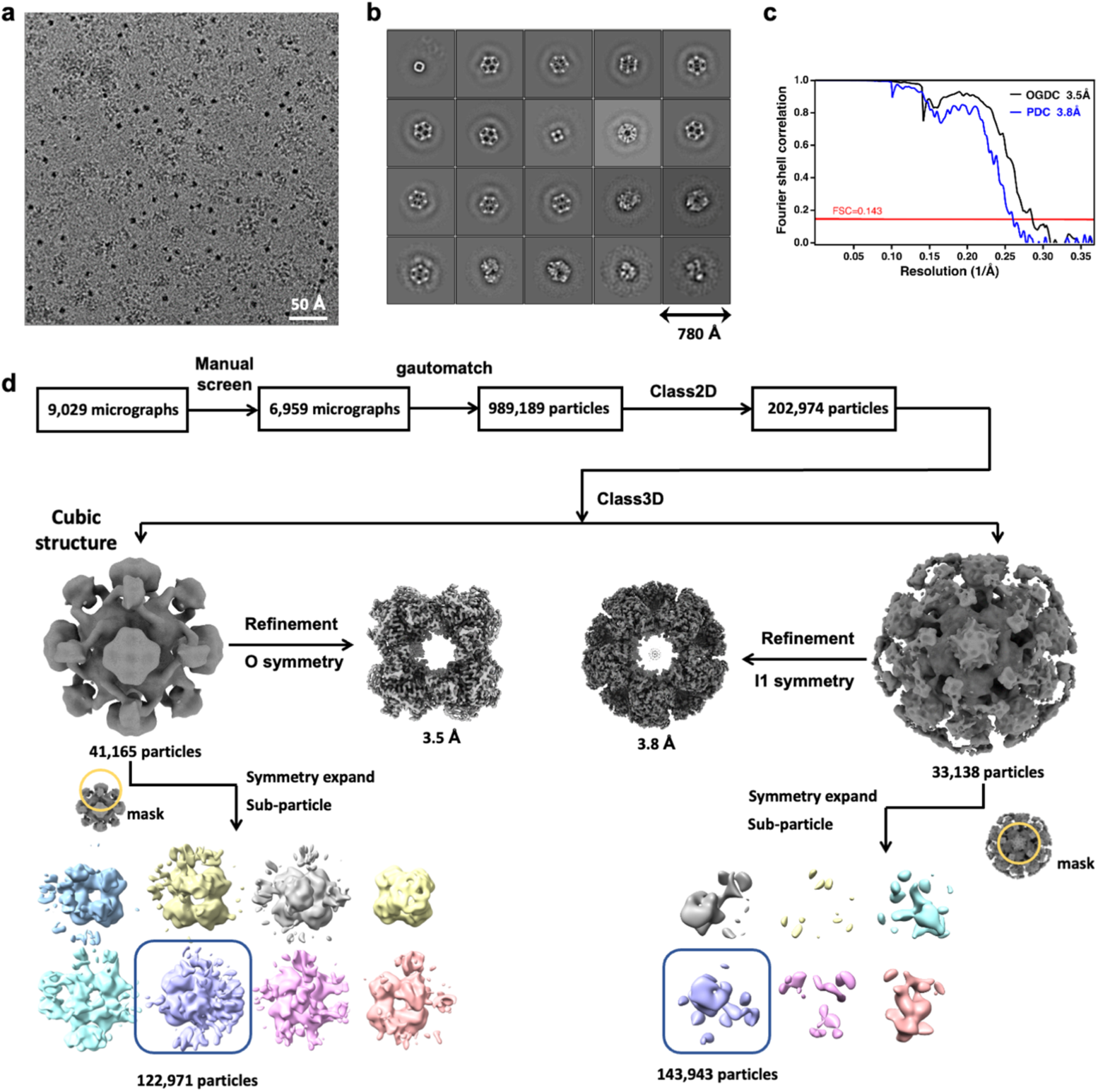
CryoEM data processing workflow. **(a)** Representative cryoEM micrograph of lysate containing PDC and OGDC. **(b)** CryoEM 2D class averages depicting icosahedral and cubic cores with smeared peripheral density. Classes of additional unidentified protein were also obtained. **(c)** FSC of E2p and E2o core. **(d)** Data processing workflow for reconstruction of E2o and E2p cores and sub-particles.

**Supplementary Fig. 4.**
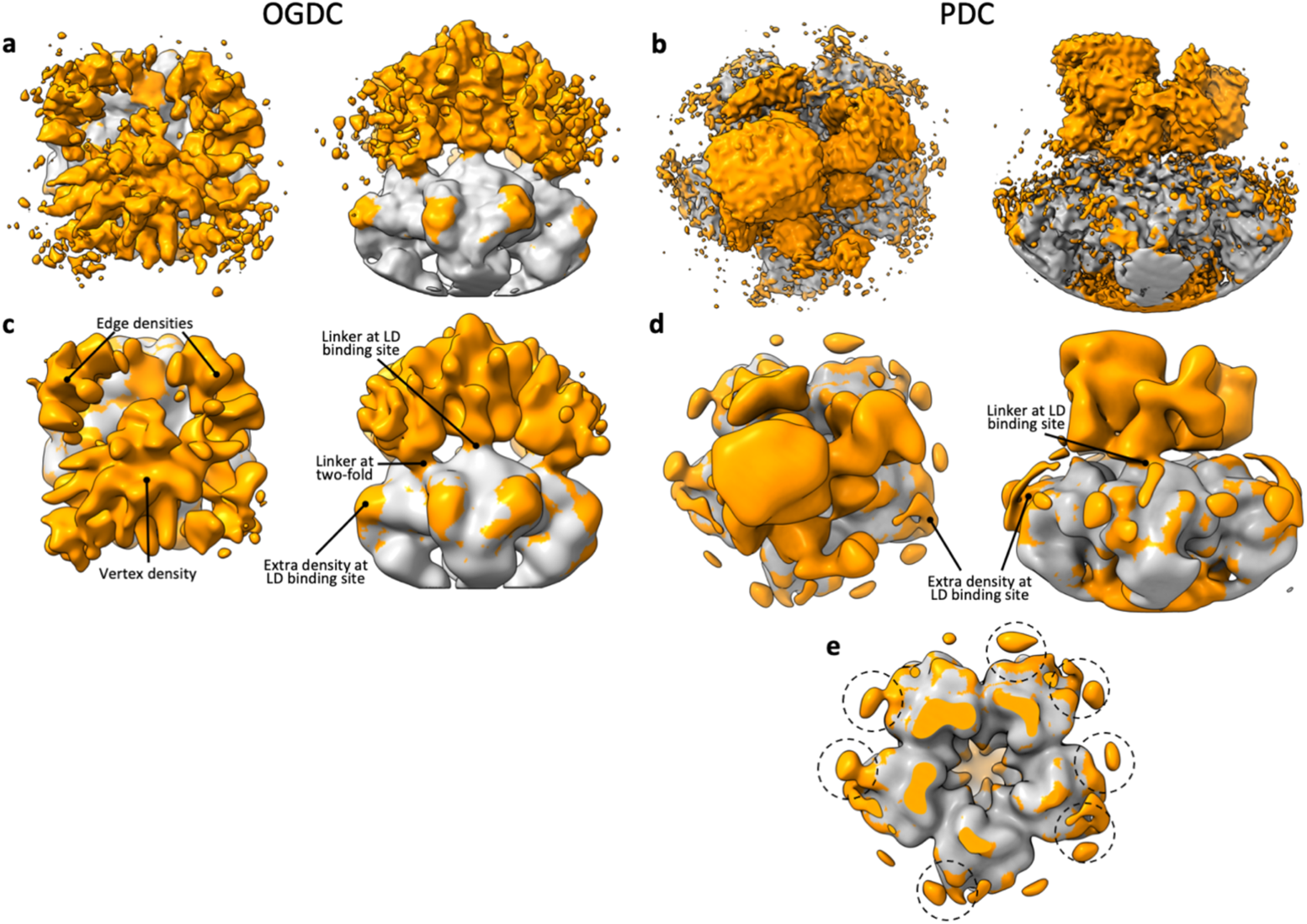
Symmetry expansion and sub-particle reconstruction of the PDC and OGDC cores. **(a)** Top (left) and side (right) views of sub-particle reconstruction with C_1_ symmetry of cubic face of OGDC. **(b)** Top (left) and side (right) views of sub-particle reconstruction with C_1_ symmetry of pentameric face of PDC. **(c)** Low-pass filtered (20 Å) OGDC sub-particle map with density external to fitted E2o model in orange. **(d)** Low-pass filtered (20 Å) PDC sub-particle map with density external to fitted E2p model in orange. **(e)** Sliced view of PDC sub-particle at LD binding site. Additional densities at LD binding site are circled.

**Supplementary Fig. 5.**
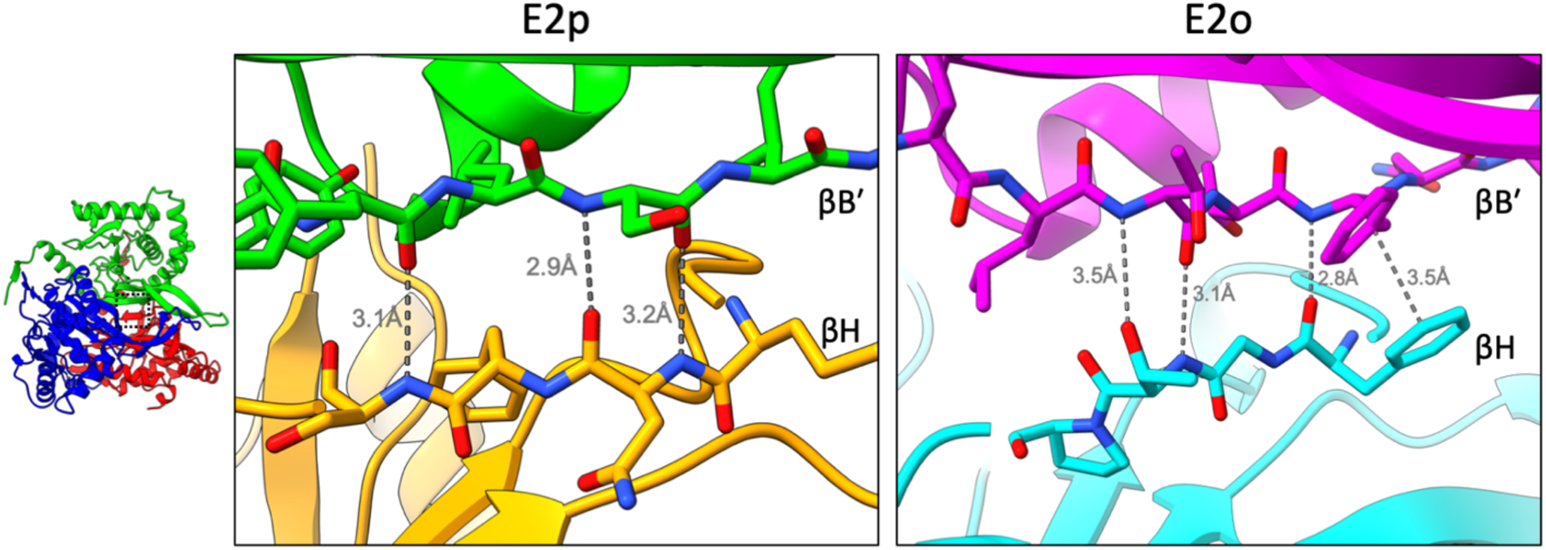
βH-βB′ interdomain interactions. View of the βH-βB′ interface within the interdomain trimer for E2p (left) and E2o (right). βH (orange; cyan) forms three hydrogen bonds with βB′ (green; magenta) in both E2p and E2o. The E2o interface has an additional slipped-parallel π-stacking interaction between Phe383 and Phe253′.

**Supplementary Fig. 6.**
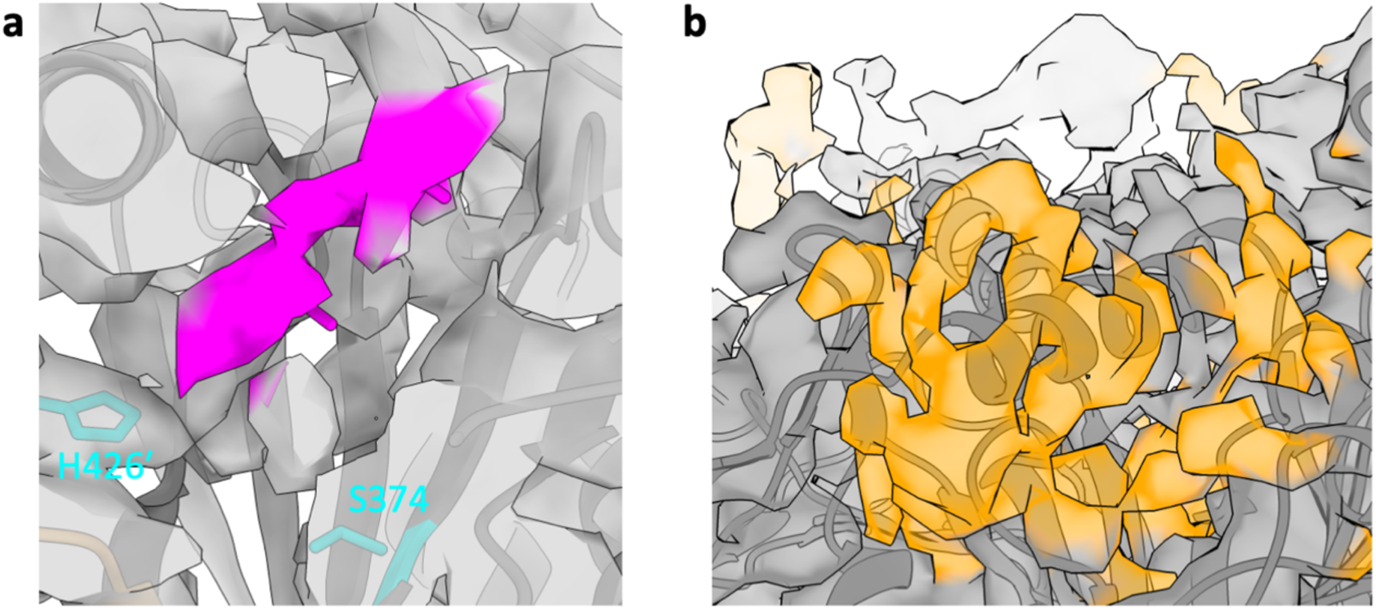
Lipoyl moiety and tentative lipoyl domain density in native E2o. **(a)** Weak, thin density within active site channel of E2o corresponding to putative lipoyl moiety (magenta). Catalytic residues (teal) indicated for reference. **(b)** Extra, cylindrical density (orange) located near previously proposed LD binding site.

**Supplementary Fig. 7.**
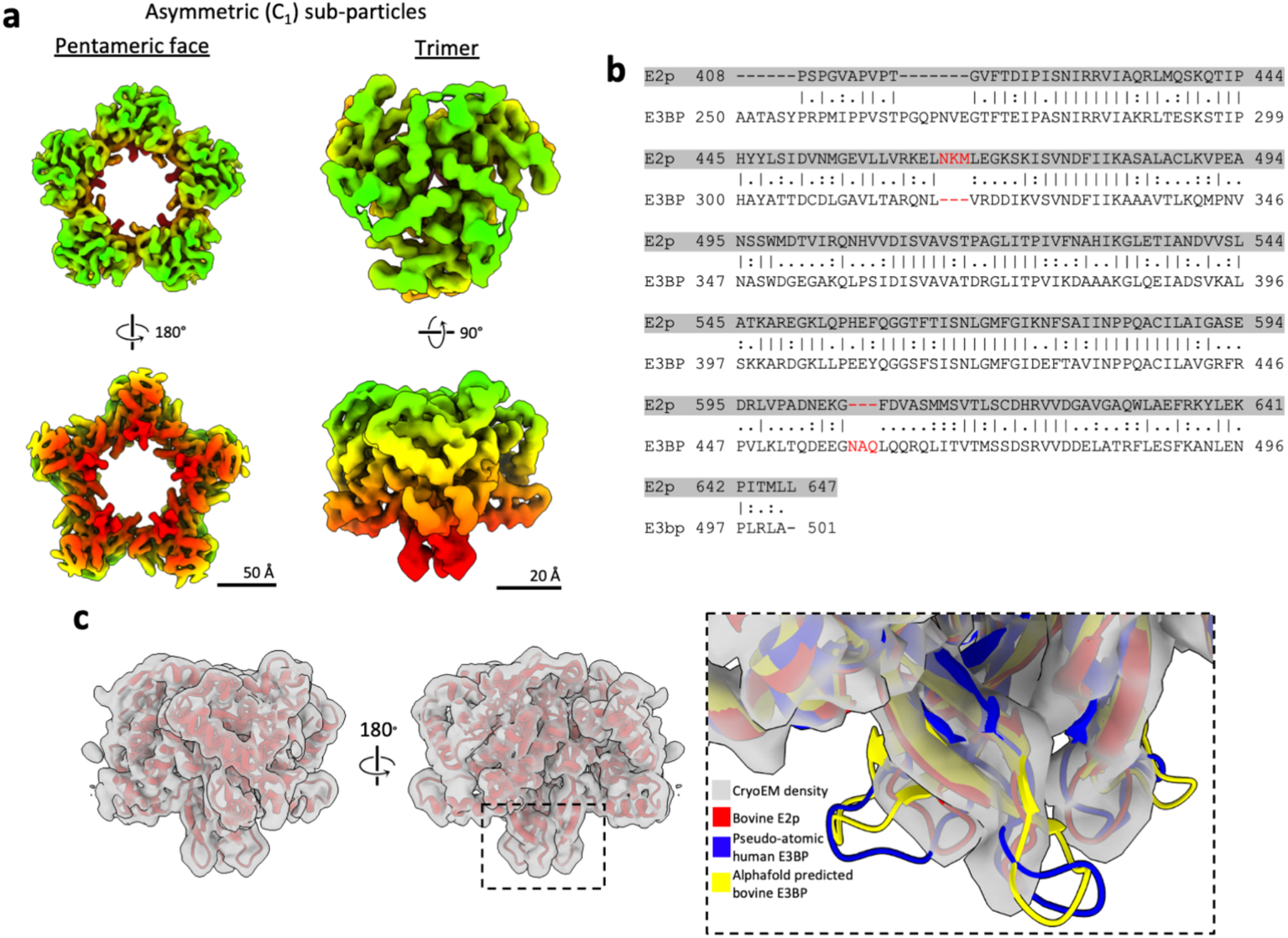
Search for E3BP in asymmetric sub-particle reconstructions. **(a)** Representative pentameric face (left) and trimer (right) sub-particle reconstructions with C_1_ symmetry. **(b)** Sequence alignment between bovine E2p and E3BP. Three-residue differences are indicated in red. **(c)** Atomic model of E2p IC domain fit into trimer sub-particle reconstruction with C_1_ symmetry. The interior loop in E3BP is extended compared to that of E2p and is a key feature for distinguishing the two components. All cryoEM densities of interior loops from each trimer class appear identical and fit E2p closely. Enlarged view of the boxed area depicts atomic models fit into cryoEM density of interior loop. The extended loop of predicted human E3BP models (PDB ID: 6H60; AF-P22439-F1) are not accommodated by the cryoEM density.

**Supplementary Fig. 8.**
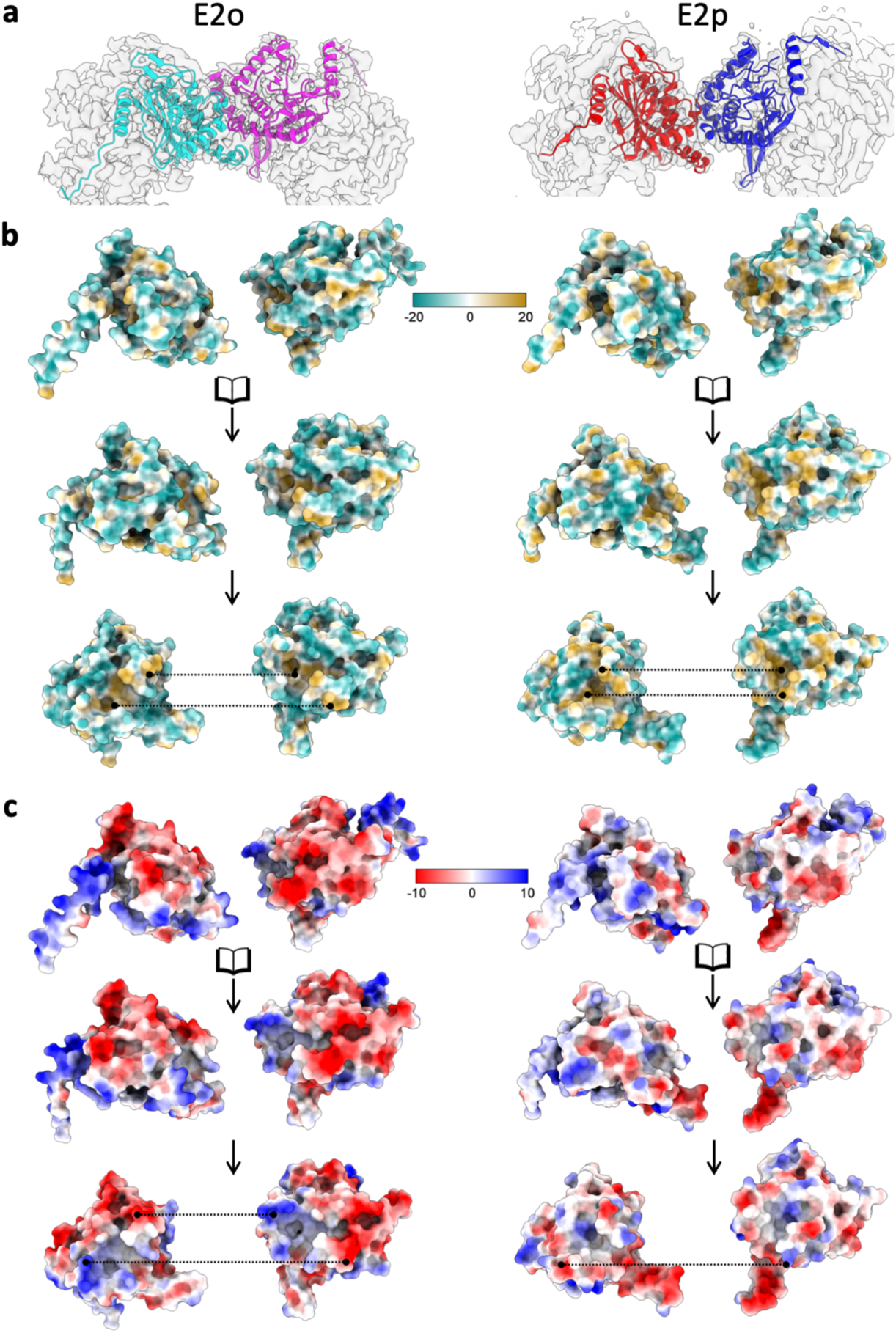
Electrostatic surface potential and hydrophobicity of E2o two-fold related intertrimer interface. **(a)** Two E2o (left) and E2p (right) IC domains at two-fold related intertrimer interface superposed in their respective density (transparent grey) and colored by chain. **(b**) Open-book view of hydrophobicity at intertrimer interface. A hydrophobic knob on each subunit binds to a complementary hydrophobic pocket on the partner subunit across the interface. Dotted lines indicate contacting surfaces. **(c)** Open-book view of electrostatic surface potential at intertrimer interface. Oppositely charged surfaces on the exterior mediate electrostatic interactions between the two subunits at the intertrimer interface.

**Supplementary Table 1.** CryoEM data collection, refinement, and validation statistics

|  | #1 OGDC<br>(EMDB-xxxx)<br>(PDB xxxx) | #2 PDC<br>(EMDB-xxxx)<br>(PDB xxxx) |
| --- | --- | --- |
| **Data collection and processing** |  |  |
| Magnification | 105,000 × | 105,000 × |
| Voltage (kV) | 300 | 300 |
| Electron exposure (e<sup>-</sup>/Å<sup>2</sup>) | 45 | 45 |
| Defocus range (μm) | -1.8 to -2.6 | -1.8 to -2.6 |
| Pixel size (Å) | 1.36 | 1.36 |
| Symmetry imposed | O | I1 |
| Initial particle images (no.) | 989,189 | 989,189 |
| Final particle images (no.) | 41,165 | 33,138 |
| Map resolution (Å) | 3.5 | 3.8 |
| FSC threshold | 0.143 | 0.143 |
| Map sharpening B factor (Å<sup>2</sup>) | -152 | -150 |
| **Refinement** |  |  |
| Model-to-map fit, mask CC | 0.88 | 0.87 |
| Model composition |  |  |
| Non-hydrogen atoms | 44496 | 104640 |
| Protein residues | 5664 | 13560 |
| Ligands | 0 | 0 |
| Mean Isotropic B factors (Å<sup>2</sup>) |  |  |
| Protein | 46.08 | 78.36 |
| Ligand | N/A | N/A |
| R.m.s. deviations |  |  |
| Bond lengths (Å) | 0.004 | 0.004 |
| Bond angles (°) | 0.944 | 0.940 |
| Validation |  |  |
| MolProbity score | 1.13 | 1.39 |
| Clashscore | 3.05 | 7.12 |
| Rotamer outliers (%) | 0.5 | 0.09 |
| Ramachandran plot |  |  |
| Favored (%) | 97.86 | 98.54 |
| Allowed (%) | 2.14 | 1.46 |
| Outliers (%) | 0.00 | 0.00 |

**Supplementary Table 2.**
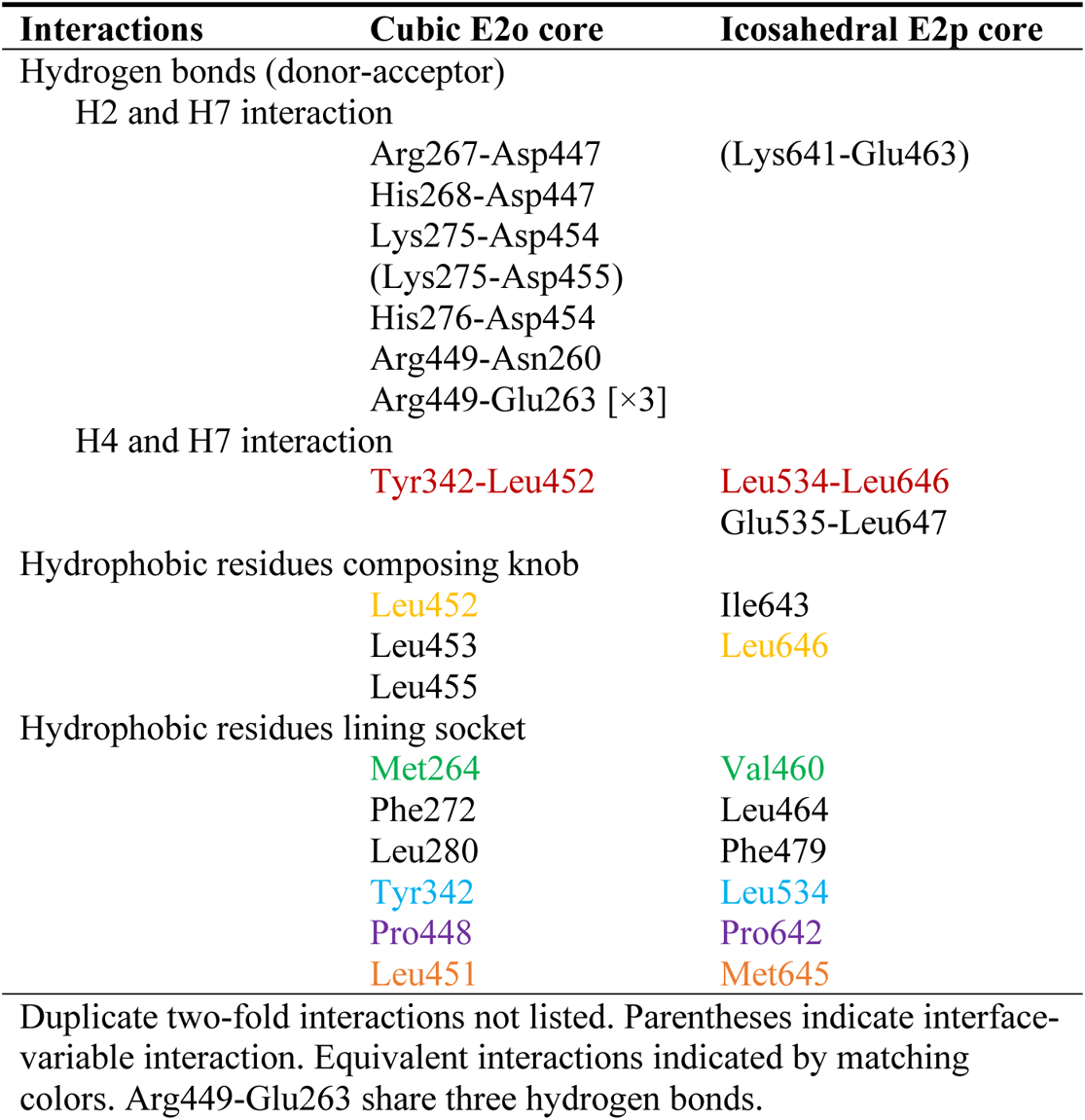
Interactions at the intertrimer interface

